# Analytical Methods and Platforms for circRNA Vector Engineering to Improve Circularization Efficiency

**DOI:** 10.1101/2024.03.25.586602

**Authors:** Yali Sun, Anis H. Khimani, Yanhong Tong, Zhi-xiang Lu

**Affiliations:** Revvity, Inc. 940 Winter Street, Waltham, MA 02451, USA

## Abstract

Circular RNAs (circRNAs) have emerged as pivotal players in RNA therapeutics. Unlike linear counterparts, circRNAs possess a closed-loop structure, conferring them with enhanced stability and resistance to degradation. This resilience makes them promising candidates for diagnostic and therapeutic applications. The ribozyme-based strategy stands out as the predominant method for synthetic circRNA production. In this strategy, ribozymes (catalytic RNA molecules) facilitate the circularization process by precisely cleaving and promoting the formation of a covalent circular structure. In the report, we detail analytical methods for circRNA vector engineering to enhance circularization efficiency. This approach will capture the attention of researchers interested in optimizing RNA circularization efficiency, as well as those focused on exploring key elements for ribozyme catalytic activity.

## Introduction

The notable success of COVID-19 mRNA vaccines has spurred increased exploration into mRNA therapeutics and related analytical methods (1–4). Circular RNAs (circRNAs), characterized by a covalently closed, single-stranded RNA (ssRNA) structure that imparts greater stability compared to their linear counterparts, hold potential as an alternative method for protein synthesis within human cells (5,6).

The circular ssRNA structures can naturally originate from mRNA back splicing, a phenomenon prevalent in eukaryotic transcriptomes that has recently garnered significant attention for its regulatory roles in diseases(5,7). However, for therapeutic applications in protein production, the development of artificial circRNAs requires specific methods (8). De novo synthesis has its limitations: the synthesized sequences are often too short, making it challenging to construct a complete protein reading frame (8,9). In vitro transcription of mRNA sequences followed by circularization using ligases is inefficient and yields substantial amounts of unwanted concatemer byproducts (10). The most widely adopted method for producing artificial circRNAs is the ribozyme approach (8,11).

The ribozyme method utilizes a permuted catalytic intron for RNA molecule self-splicing. This permuted intron-exon (PIE) consists of fused partial exons (E1, E2 in ***Figure 1A***) flanked by half-intron sequences (scribble lines/hairpin structures in ***Figure 1A***) (12). The reaction involves the use of GTP and Mg^2+^ as cofactors and entails two transesterification steps at the 5’ and 3’ splice sites for intramolecular end joining (***Figure 1B***). In comparison to chemical and enzymatic ligation, the ribozyme method is more suitable for cyclizing longer linear RNA precursors, and its reaction conditions and purification methods are simpler (11).

**Figure 1:**
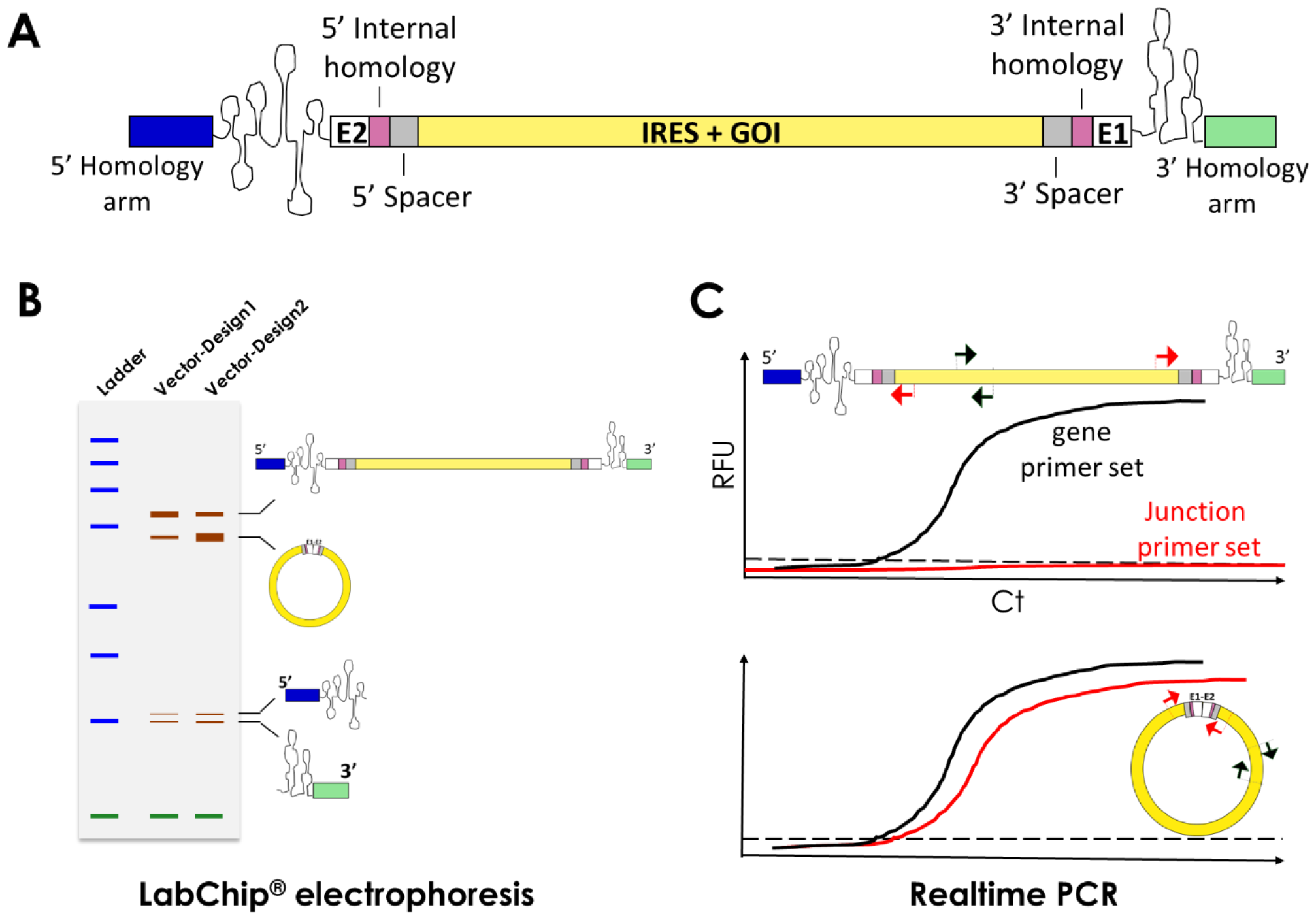
Analytical analysis workflow of synthetic RNA circularization. **(A)** A typical permuted intron-exon construct design for synthetic circRNA precursor. E1, E2: exon 1, exon2; IRES: internal Ribosome entry site; GOI: Gene-of-Interest; Catalytic intron as a preserved element critical for ribozyme folding is drawn as scribble lines and hairpin structures. **(B)** Schematic diagram showing fragment analysis of RNA species in RNA circularization on LabChip®. Construct variants by circRNA vector engineering are labeled as Vector-Design1 and Vector-Design2. **(C)** Real-time PCR primer design and amplification curve for measuring ribozyme circularization. Top diagram represents results from linear precursor or samples with no or low circulation yield; bottom diagram represents results from samples with success circularization. Red arrows represent junction primer sets; black arrows represent body primer set.

Through rational design of precursor PIE construct by Wesselhoeft et.al (13), homology arms are strategically positioned at 5’, 3’ ends of the precursors with the aim of aligning splice sites into proximity of one another (blue, purple, and green boxes in ***Figure 1A***); The spacers (gray box in ***Figure 1A***) are also intentionally inserted before internal ribosome entry site (IRES) and after the coding sequence (CDS) with the aim of reducing IRES interfering ribozyme folding. The design principles outlined above are driven by a singular objective: optimizing circRNA backbones for enhanced RNA circularization. Consequently, systematic high-throughput analytical techniques play an increasingly vital role in evaluating the circularization of RNA, particularly in the field of circRNA ribozyme backbone engineering.

In this study, we undertook the process development of synthetic circRNA using ribozyme method. We demonstrate a high-throughput analytical solution for assessing the efficiency of synthetic RNA circularization, employing the Labchip® GX Touch™ electrophoresis platform and real-time PCR relative quantification platform (**Figure 1B & 1C**). These approaches will facilitate the discovery process for researchers aiming to enhance RNA circularization efficiency, as well as those delving into the exploration of crucial elements influencing ribozyme catalytic activity.

## Material and Methods

### CircRNA vector design and mutagenesis

T4 phage and Anabaena permuted intron-exon (PIE) were used as circRNA backbones (***Table 1***). Other elements such as homology arms, internal homology, spacers were designed following a previous study, gene of interests with variable sizes were inserted between exon fragments (13) (***Figure 1A***). Synthesis of these fragments and their insertion into the pUC19 backbone were accomplished through gene synthesis services provided by Genscript. Construct_1 vector incorporated the Gluc T4PIE EMCV sequence, Construct_2 vector utilized CVB3-Gluc-pAC sequence, and Contruct_3 was derived from Construct_2 vector by removing CVB3 IRES region. To access the complete sequences of Gluc T4PIE EMCV and CVB3-Gluc-PAC, kindly refer to the previous study (13). Site-directed mutagenesis constructs (i.e. Construct_2_mut, Construct_3_mut) was performed by the Megaprimer method (14), with details of primers in ***Table 2***.

**Table 1:**
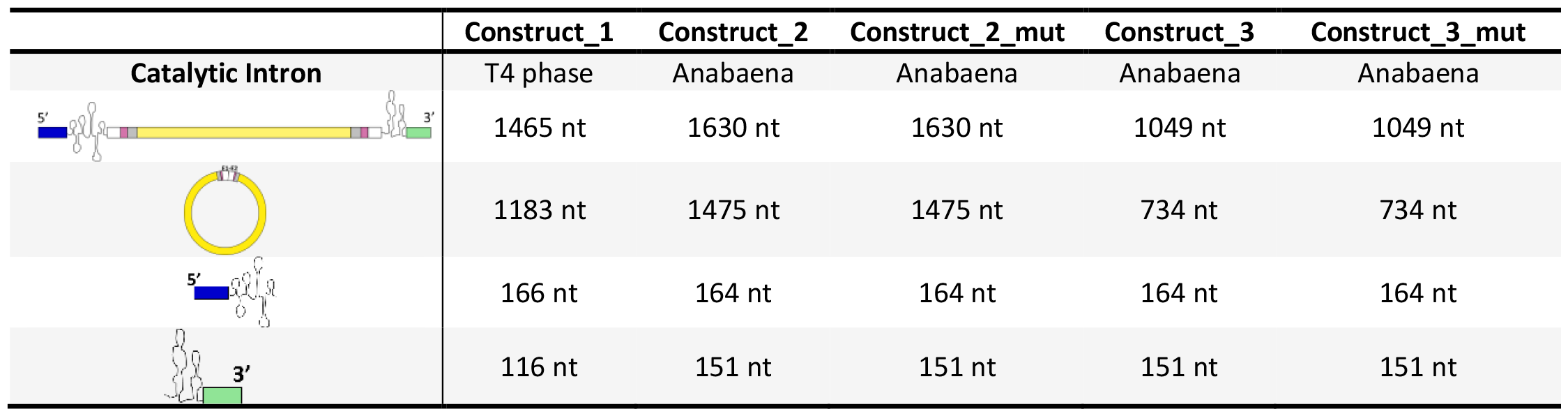
Sequence features of circRNA plasmids.

|  | Construct_1 | Construct_2 | Construct_2_mut | Construct_3 | Construct_3_mut |
| --- | --- | --- | --- | --- | --- |
| **Catalytic Intron** | T4 phase | Anabaena | Anabaena | Anabaena | Anabaena |
|  | 1465 nt | 1630 nt | 1630 nt | 1049 nt | 1049 nt |
|  | 1183 nt | 1475 nt | 1475 nt | 734 nt | 734 nt |
|  | 166 nt | 164 nt | 164 nt | 164 nt | 164 nt |
|  | 116 nt | 151 nt | 151 nt | 151 nt | 151 nt |

**Table 2:**
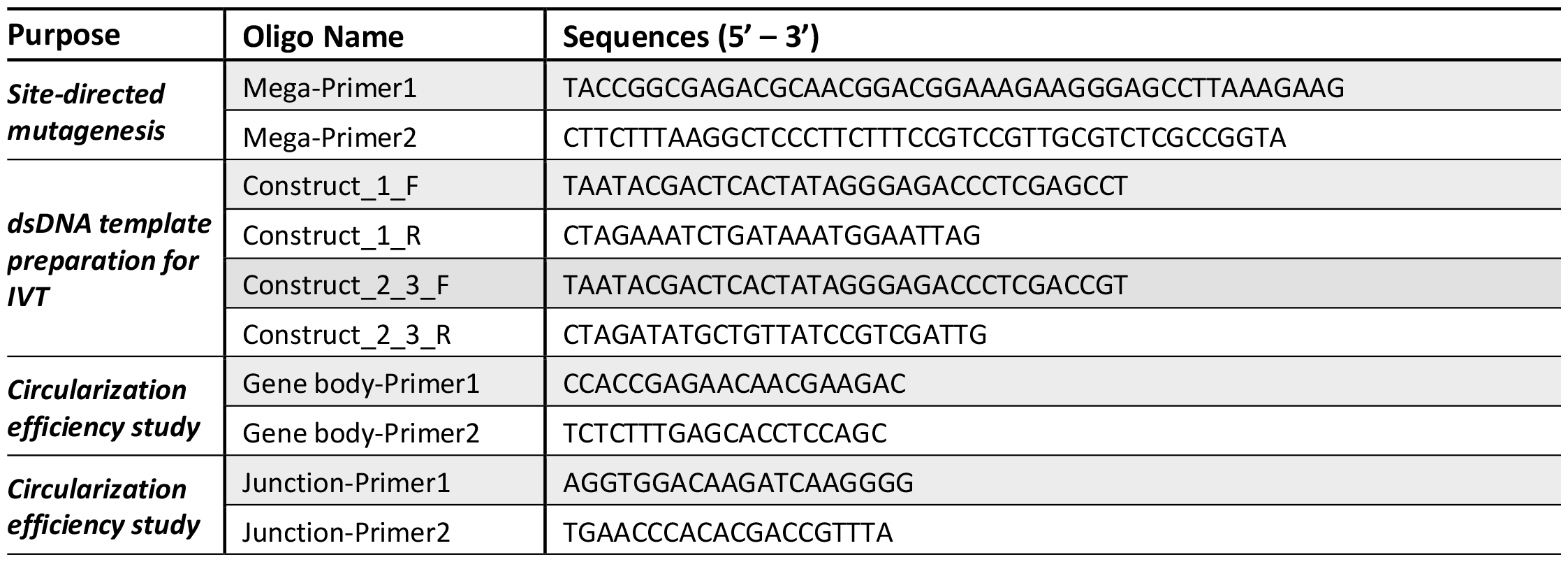
Sequences of primers.

### CircRNA synthesis

T7 double stranded DNAs (T7-dsDNAs) were amplified by T7-tagged primer pairs (***Table 2***) by Phire Hot Start II PCR Master Mix kit (Thermo Fisher, Cat # F125L) using proper plasmids as templates, and purified by Zymo oligo clean and concentrator kit (Zymo, cat # D4060). RNAs were synthesized by in-vitro transcription (IVT) at 37° C for 2 hrs from 1 µg purified dsDNAs using a T7 high yield RNA synthesis kit (NEB, Cat # E2040S) in a total volume of 20 µL. After IVT, 20 µL of reactions were treated with 2 µL DNase I (NEB, Cat # M0303S) in a total volume of 100 µL at 37°C for ∼ 20 min before further analysis and treatment.

### LabChip® electrophoresis

Concentration of RNA products from IVT were measured by Nanodrop. All RNA samples concentrations were adjusted to 1 ng/µL for optimal performance using the LabChip® RNA Pico Assay (Part # CLS960012). Samples at 6 μl volume were transferred to 96-well plate, subsequently heated to 70° C for 2 min, and immediately placed on ice for 3 min. Then the 6 µL of samples were mixed with 24 µL 1x sample buffer (provided in LabChip® RNA Pico Assay) and loaded on the LabChip® GX Touch™ (Part # CLS137031) with RNA Pico reagent, using the DNA 5K/RNA/Charge Variant Assay Chip (Part # 760435). HT RNA Pico sensitivity program was selected for the LabChip® run (***Figure 2B***). Peak height or Peak area given by LabChip® GX Reviewer software was used for data analysis.

**Figure 2:**
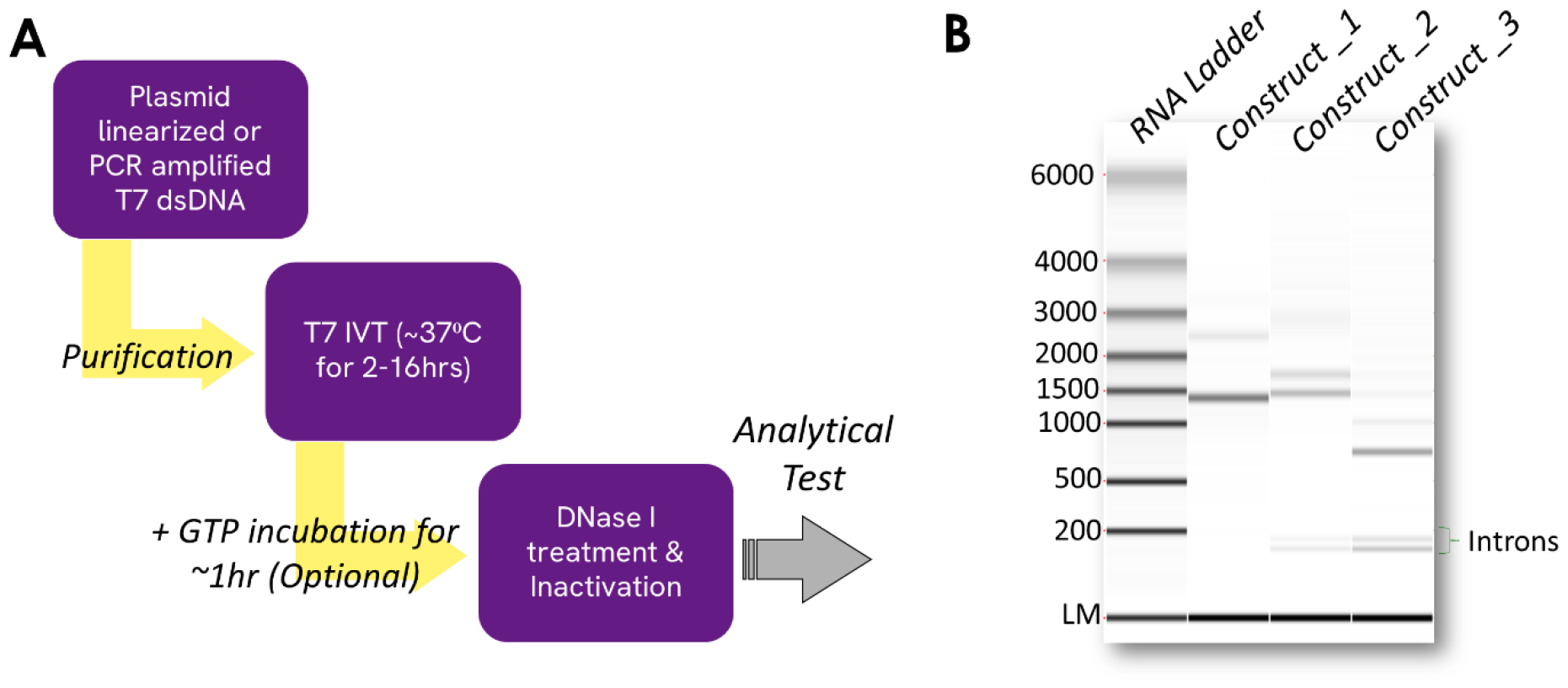
In-Vitro Transcription (IVT) and circularization. **(A)** Workflow of ribozyme-based RNA circularization procedures. **(B)** LabChip® gel confirmation of precursor RNA circularization. Construct_1 is not subjected to circularization process, only shows precursor RNA band representing the large molecular size. Construct_2 and Construct_3 are subjected to circularization process, and two intron ends are spliced out as two small size bands (< 200 nt).

### Reverse transcription and real-time PCR

Universal primers were designed at gene-body region (labeled as gene primer set) and at splicing-ligation junction region (labeled as junction primer set) in ***Figure 1C***. A total of 1E5 copy RNA template was reverse transcribed and real-time PCR amplified by reagent A and enzyme mix from Revvity Respiratory SARS-CoV-2 RT-PCR Panel 1 reagent kit (part # SDX-56390), together with EvaGreen (Biotium, Cat # 31000) using either gene primer set or junction primer set with final concentration of primers at 333 nM. Real-time PCR cycles were run as 37° C/2 min., 50° C /15 min., 94° C/10 min., 40 cycles (94° C/10s, 60° C/10s, 65° C/45s) and melting curve was analyzed at 30° C to 95° C temperature range. For each sample, the real-time PCR was performed in triplicate on QuantStudio™ real-time PCR system. Averages of the ***Ct*** values were processed for further analysis. Primers used in PCR are listed in ***Table 2***.

### RNase R cleanup linear RNA

Total 500 ng of non-purified circRNAs were used for RNase R treatment (Lucigen, Cat # RNR07250). The reaction was performed at 37° C for 30 min with 1 x RNase R buffer and 1 µL (“+”) or 0 (“-”) µL RNase R in a 20 µL volume. After RNase R treatment, the samples were diluted to 1 ng/µL by water, heated to 70° C for 2 min, and immediately placed on ice for 3 min for subsequent LabChip® electrophoresis analysis as described above.

## Results

### A model for evaluating the production of synthetic circRNA

Two transesterification reactions take place in the ribozyme-based circularization process (13), resulting in circularization of the intervening region and excision of 5’ intron and 3’ intron (***Figure 1A*** and ***1B***). Compared to the precursor RNA molecules which are in linear form, the circularization reaction would produce a circular RNA with a shorter length, and excised intron products (13). We aimed to explore the capability of LabChip® electrophoresis system to differentiate circularized RNA products from other RNA species in circularization reaction. If successful, this could potentially enable the high throughput analytical analysis of circularization efficiency (***Figure 1B***). Likewise, in the intervening region (yellow box in ***Figure 1C***), by designing real-time PCR gene primers which target both precursor and circRNAs, and junction primers which only amplify when circularization occurs (***Figure 1C***). Thus, researchers can effortlessly quantify the relative circularization efficiency in circRNA vector engineering by coupling high throughput electrophoresis and qPCR test.

### Excised introns signaling the occurrence of circularization

To test the aforementioned analytical model, we conducted in vitro transcription and circularization using three circRNA constructs: Construct_1 is T4 page PIE, while Construct_2 and construct_3 use Anabaena PIE with different sizes of intervening region (***Table 1***). After the IVT reaction and DNase I treatment, all three samples were diluted to 1 ng/µL in water for electrophoresis on the LabChip® platform. As shown in ***Figure 2B***, Construct_1 has one main band at ∼1400 nt, which is the expected size of its linear RNA precursor fragment. We didn’t see any excised intron fragments indicating that RNA circularization did not occur or had a very low efficiency of circularization (***Figure 2B***). In contrast, Contruct_2 and Construct_3 which use Anabaena self-splicing intron backbone show two small fragments under 200 nt. They are more likely excised 5’ and 3’ intron fragments (***Figure 2B*** *and* ***Table 1***). These small fragments serve as a good indicator that the circularization process is occurring in Construct_2 and Construct_3. Given that we load an equal amount of total RNA, an increased peak signal of 5’, 3’ intron fragments in Construct_3 indicates a higher efficiency in circularization. These observations agree with prior studies showing the shorter length of “IRES+GOI” region, the higher circulation efficiency with the same circRNA PIE backbone(13).

In addition to the small fragments, the two large peaks in Construct_2 and Construct_3 are more likely related to circRNA forms and linear precursor RNAs. Earlier investigations have demonstrated that the migration behavior of circRNA towards larger precursor RNAs or equivalently weighted nicked circRNAs is largely influenced by the electrophoresis system employed (15) . Subsequent experiments were devised to comprehensively characterize these two prominent peaks.

### Peak characterizations of linear and circular RNAs

Since the splice site in E1, E2 intron boundary is a key region for self-splicing, we have reasons to believe that by disrupting this region, the circularization efficiency will decrease considerably resulting in the reduction of the circRNA form upon electrophoresis. To investigate this, we generated two additional constructs, namely Construct_2_mut and Construct_3_mut, incorporating 5’ splice site mutations in the E1 splice site. Following in vitro transcription (IVT) and circularization, the samples were analyzed on the LabChip® platform alongside their respective references, Construct_2 and Construct_3. As anticipated, the intensity of the small intron fragments noticeably diminished and became scarcely detectable, indicating a reduction in circularization efficiency in these mutant constructs (***Figure 3A***). In Construct_2_mut and Construct_3_mut, the two larger fragments, exhibiting weakened peaks with relative smaller sizes in comparison to their respective references, are hypothesized to represent the circRNA forms (***Figure 3A***).

**Figure 3:**
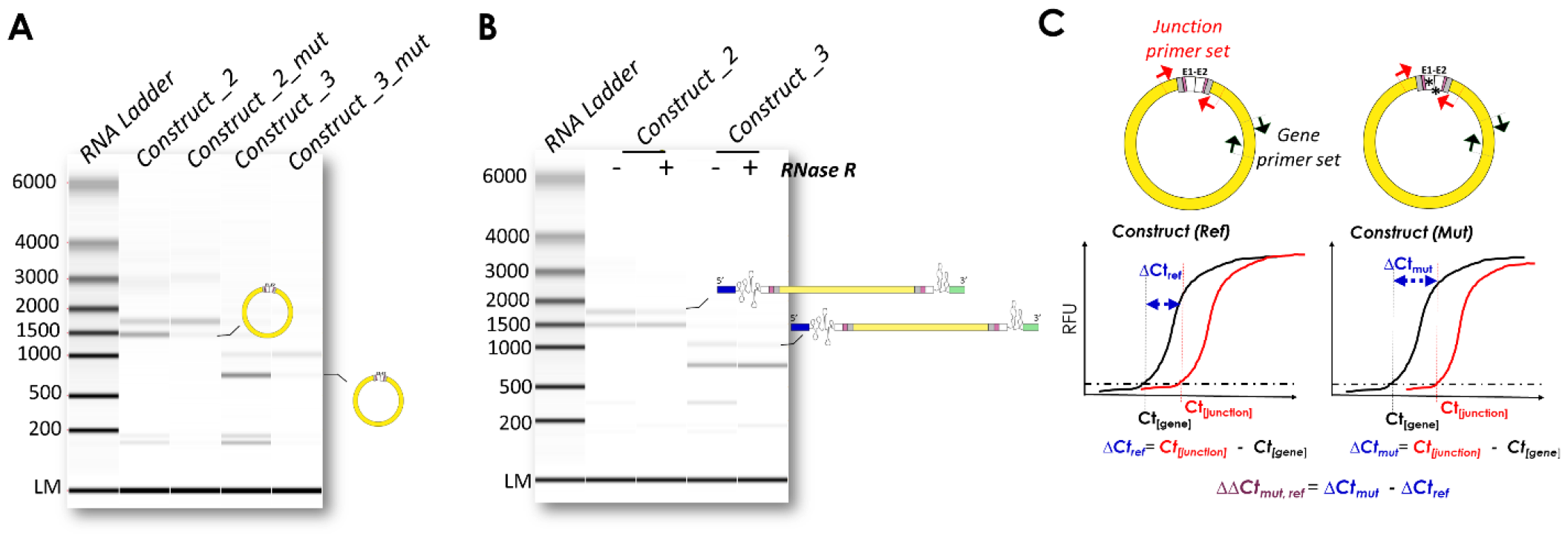
Peak characterization and analytical test of circRNAs. **(A)** Splice site mutations (Construct_2_mut and Construct_3_mut) in circRNA vectors reduce circularization efficiency. **(B)** CircRNAs are enriched by RNase R treatment. **(C)** A relative circularization efficiency calculation by real-time PCR between reference construct and mutant construct. ΔΔCt_mut, ref_ <0 means mutant construct improves circularization efficiency; ΔΔCt_mut, ref_ >0 means mutant construct decreases circularization efficiency. ΔΔCt_mut, ref_ =0 means mutant construct doesn’t impact circularization efficiency. *: arbitrary mutations.

To confirm that the upper peak corresponding to circRNA peak is the linear RNA precursor, we subsequently subjected Construct_2 and Construct_3 RNA samples to treatment with RNase R, an enzyme that specifically digests linear RNA, thus, employed for circRNA cleanup. If the upper peak indeed signifies the linear RNA precursor, a significant reduction would be expected in its intensity following RNase R treatment, while the circRNA peak would remain unchanged. As illustrated in ***Figure 3B***, after RNase R treatment, the upper peaks (approximately 1.7 kb for Construct_2 and 1 kb for Construct_3) exhibited decreased intensity, validating their identity as linear RNA precursors, whereas the circRNA peaks (around 1.5 kb for Construct_2 and 750 nt for Construct_3) remained unaffected.

### Circularization efficiency estimation of circRNA constructs

With the confirmed identification of the circRNA peak and linear RNA precursor peak, we then calculated the circularization efficiency for each construct using ***Peak Area Ratio***: *circRNA / (circRNA + linear RNA) * 100%*. In ***Figure 3A***, Construct_2 exhibits a circularization efficiency of 61.6% compared to Construct_2_mut, which has a circularization efficiency of 14.7%. Similarly, Construct_3 demonstrates a circularization efficiency of 86.1%, while Construct_3_mut exhibits a circularization efficiency of 18.5%.

As an orthogonal method confirmation of our observations above, relative quantification by realtime qPCR was also performed on these samples using both the gene primer set and junction primer set. As depicted in ***Figure 1C*** and ***3C, Ct***_***[gene]***_ is utilized to measure both precursors and circRNAs, while ***Ct***_***[junction]***_ is specifically employed for cirRNA measurement. The bigger ***Ct*** value means the lower amount of target molecules. In theory, ***ΔCt (Ct***_***[junction]***_***-Ct***_***[gene]***_) could serve as a direct measure of circularization efficiency. However, owing to amplification efficiency bias among primer sets, without adjustment or calibration, the absolute ***ΔCt (Ct***_***[junction]***_***-Ct***_***[gene]***_) lacks precision. In this context, we applied relative quantification method ***ΔΔCt***_***mut***,***ref***._ (= ***ΔCt***_***mut***_ ***– ΔCt***_***ref***_) for data analysis. Given that our primers are designed in common regions of these circRNA constructs, ***ΔΔCt***_***mut***,***ref***_ emerges as a valuable solution for conducting relative circularization efficiency analysis, which will allow us to discern the circRNA with the optimal circularization efficiency. If a mutant construct reduces circularization efficiency compared to reference construct, ***Ct***_***[junction]***_ will be delayed, ***ΔCt (Ct***_***[junction]***_***-Ct***_***[gene]***_) of mutant construct will become larger, then ***ΔΔCt***_***mut***,***ref***_ will be >0. For instances, ***ΔCt*** (construct_2) as reference is 4.4, ***ΔCt*** (construct_2_mut) is 6.4, ***ΔΔCt***_***mut***,***ref***_ *of contruct_2_mut is 2;* ***ΔCt*** (construct_3) as reference is 4.8, ***ΔCt*** (construct_3_mut) is 6.4, ***ΔΔCt***_***mut***,***ref***_ *of contruct_3_mut is 1*.*6*. Both positive scores of ***ΔΔCt*** indicate circularization efficiency decrease in construct_2_mut and construct_3_mut, which is an independent result aligning very well with LabChip® electrophoresis fragment analysis.

## Conclusion and Discussion

We characterized peaks from circRNA via processing unpurified circRNA samples in LabChip® electrophoresis (***Figure 4A***). The LabChip® RNA Pico assay exhibits remarkable performance in differentiating excised introns, circRNA, and precursor RNA. It is important to note that this performance of the LabChip® RNA Pico assay may vary depending on sample composition (RNA sequence design, secondary structure, nucleotides components, etc.) (16). Labchip® RNA Pico assay, utilizing size separation, offers valuable information on circularization process. However, to capture all aspects of final functional products, an orthogonal approach such as a real-time PCR platform is beneficial. For instance, in ***Figure 4A***, the circRNA form (theoretical size = 1475 nt, predicted size based on LabChip® linear RNA Ladder = 1466 nt) may not be easily distinguishable from its equal-weight nicked circRNAs, as the migration behavior of circRNA in Labchip® RNA Pico assay closely mirrors that of its linear-sized RNA molecule. In some cases, trace amount of fragments with larger sizes than the two peaks (circRNA and linear precursor RNA) were observed, the potential structures of which are under investigation. Real-time PCR platform, employing junction primer design (***Figure 1C***), facilitates the quantification of ligated products. Nevertheless, it is unable to differentiate between intra-circularization and concatemer ligation by products. In our study, we aimed to emphasize the complementary nature of Labchip® electrophoresis fragment analysis and real-time PCR platforms in providing a comprehensive understanding of the circRNA bioprocess development (***Figure 4B***).

**Figure 4:**
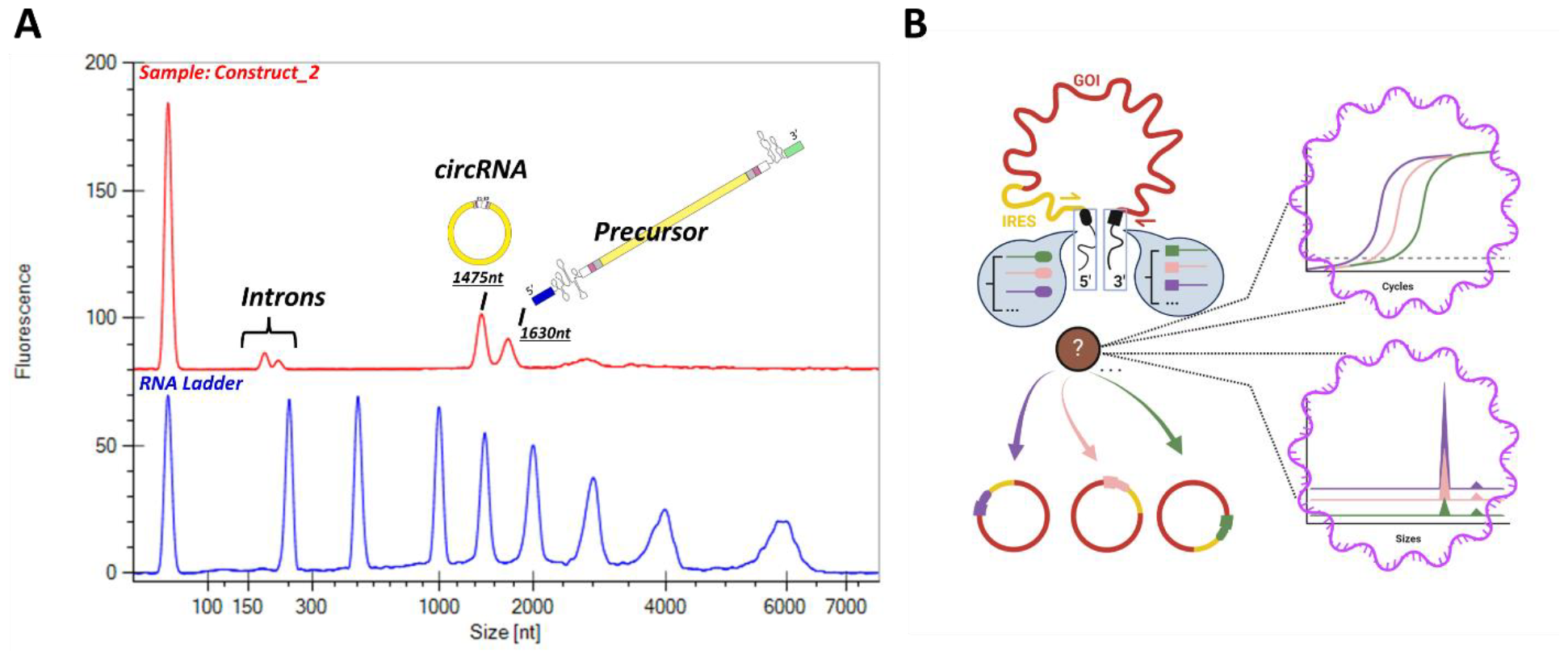
Analytical solution for the evaluation of RNA circularization efficiency. **(A)** circRNA peak patterns using RNA pico reagents on LabChip®. Top panel: One typical non purified RNA product by ribozyme-based circularization. Bottom panel: RNA Ladder from RNA Pico reagent. If size resolution of electrophoresis platform is optimal, assuming no 100% circularization efficiency, four bands are usually expected: small introns’ peaks excised from 5’ end and 3’ end, one circRNA peak, and one RNA precursor peak. The sizes labeled below RNA species are theoretical value based on sequence design. **(B)** Graphic illustration of RNA circularization efficiency evaluation using LabChip® platform and qPCR platform. This illustration was created with BioRender.com.

The analytical methods employed to test circRNA circularization efficiency hold significant implications. By enhancing the ability to engineer circRNA vectors with improved circularization efficiency, this approach not only enables researchers aiming for circRNA construct backbone with higher circularization formation, but also facilitates the study of RNA structures and sequences in ribozyme function. We believe our analytical methods not only refine circRNA upstream process development but also fuel a broader understanding of RNA biology, positioning itself as a valuable tool for both applied and fundamental research in the fields of biotechnology and molecular biology.

## Funding

Revvity, Inc. provided all the funding for this research.

## Conflict of Interest Statement

Although the authors are employed and funded by Revvity, Inc., this should not detract from the objectivity of data generation or interpretation.

## Acknowledgements

Z.L. conceived and designed the study. Z.L. and Y.T. supervised the project. Y.S. performed experiments. Z.L. and Y.S. carried out the analysis and drafted the paper, A.K provided critical insights. All authors reviewed the paper.

## Data availability statement

The data underlying this article are available in the article or upon request.

